# Comprehensive investigation of sleep architecture by *APOE* genotype: *APOE4* homozygotes have higher arousal thresholds

**DOI:** 10.1101/2025.05.30.656615

**Authors:** Gawon Cho, Anne Chen, Eunyoung Choi, Orfeu M. Buxton, Daniel Kay, Brienne Miner

## Abstract

**Background and Objectives:** *APOE4*, a genetic risk factor for Alzheimer’s Disease (AD), is associated with reduced functional connectivity of brain regions that regulate sleep, which may predispose persons to AD via altering sleep architecture. However, little is known about differences in sleep architecture by *APOE* genotype.

**Methods:** This cross-sectional study examined the association between *APOE* genotype and sleep architecture among middle-aged and older adults, using polysomnography (Sleep Heart Health Study, N=3,132). *APOE* genotype included: *APOE4* heterozygotes, *APOE4* homozygotes, *APOE2* carriers, and *APOE3* homozygotes. Macro sleep architecture was quantified using the percentage of time spent in rapid eye movement sleep (%REM), N1 (N1%), N2 (%N2), N3 (%N3), and arousal index. Micro sleep architecture was quantified as odds ratio product (ORP; a continuous measure of sleep depth) for overall sleep and each sleep stage, and spindle characteristics (power, density, and frequency). Linear regression was used, adjusting for covariates.

**Results:** The mean age was 67, 53 percent were female, 24% were *APOE4* heterozygotes, 2% were *APOE4* homozygotes, 14% were *APOE2* carriers, and 60% were *APOE3* homozygotes. Macro sleep architecture did not vary across genotypes. Compared with *APOE3* homozygotes, *APOE4* homozygotes exhibited fewer arousals with age (β=-0.33 per hour/year, p=0.04), resulting in significantly fewer arousals at age 70+. ORP decreased in a dose-response pattern with the number of *APOE4* alleles during overall sleep and across all sleep stages (ORP_APOE3/3_=0.94, ORP_APOE3/4_=0.91, ORP_APOE4/4_=0.87), and these group difference widened with each year of age. Finally, there was a trend for lower spindle density and power in *APOE4* homozygotes relative to *APOE3* homozygotes (*p*s=0.06).

**Conclusions:** Arousal threshold increased in a dose-response manner with each *APOE4* allele, as evidenced by the findings on ORP. The differences in ORP between *APOE3* homozygotes and *APOE4* carriers widened further with age, paralleling age-related declines in arousal index among *APOE4* homozygotes. Despite these indications of elevated arousal thresholds that might suggest less sleep fragmentation in *APOE4* carriers, *APOE4* homozygotes exhibited poorer sleep micro architecture, including trends toward reduced sleep spindle activity. Taken together, reduced arousability in *APOE4* carriers may reflect abnormalities in cortical activation that blunt arousal rather than an indicator of healthier sleep.

## 1. INTRODUCTION

The *apolipoprotein E ε4* (*APOE4*) is the most established genetic risk factor for Alzheimer’s disease (AD), a progressive neurodegenerative disorder affecting more than six million older adults in the U.S. ^1^ *APOE*4 carriers, including heterozygotes and homozygotes, exhibit reduced functional connectivity of key neuromodulatory regions that regulate sleep, including the basal forebrain^2^ and the locus coeruleus. ^3^ These regions are central to regulating the sleep-stage transitions that define sleep architecture, which refers to the organization of sleep period into stages of varying depth, including rapid eye movement (REM) sleep and non-REM stages N1, N2, and N3 (slow-wave sleep; SWS), as well as arousal events. ^4–8^ Furthermore, locus coeruleus is the primary source of norepinephrine in the central nervous system, playing a key role in the generation of sleep spindles, ^9^ i.e., isolated bursts of rhythmic brain activity that typically occurs during N2 and contributes to memory consolidation. ^10–12^ Disrupted connectivity among brain regions that regulate sleep in *APOE4* carriers may increase vulnerability to alterations in sleep architecture, such as sleep fragmentation, a known risk factor for AD and related neurodegeneration. ^12–15^ Thus, impaired sleep architecture may be a mechanistic pathway linking *APOE4* genotype to elevated AD risk.

To date, no study has examined differences in sleep micro architecture by *APOE* genotype. Only two studies have evaluated differences in sleep macro architecture by *APOE4* genotype, and they have reported heterogeneous findings. ^16,17^ Among older men, Tranah et al. ^16^ found *APOE*4 carriers to spend more time in SWS, whereas the investigation among among older men and women by André et al. ^17^ found *APOE4* carriers to have comparable SWS but reduced REM sleep. Both studies relied on relatively small samples with limited representation of *APOE4* carriers, particularly homozygotes, which could have restricted statistical power to detect robust genotype-specific sleep characteristics. Moreover, both studies examined adults in late adulthood, a stage in life where sleep disturbances are highly prevalent regardless of genetic risk, making the detection of genotype-related differences more difficult. ^18–20^ In addition, only one of the two studies included women, despite their higher susceptibility to both AD and various forms of sleep disruptions. ^21–24^ Notably, *APOE4* appears to confer greater AD risk in women, underscoring the importance of characterizing sleep architecture in a sample inclusive of women.

To address these gaps, the present study examined associations between *APOE* genotype and both sleep macro- and micro-architecture in a large, community-based sample of middle-aged and older adults, with robust representation of *APOE4* heterozygotes and homozygotes. Based on prior evidence showing reduced functional connectivity in sleep-regulating brain regions among *APOE4* carriers, ^4–8^ we hypothesized that they would exhibit altered sleep architecture characterized by lighter sleep depth (for example, lower SWS and more frequent arousals), reduced spindle activity, and reduced REM sleep relative to noncarriers. We further hypothesized that genotype-associated differences would be more pronounced in middle age than in late adulthood, given the high prevalence of sleep disturbances in later life regardless of genetic risk. Finally, motivated by prior findings that *APOE4* confers greater AD risk in women, we explored whether the associations between *APOE4* genotype and sleep architecture differed between females and males. These investigations represent an essential first step in assessing whether alterations in sleep architecture may serve as a modifiable risk factor for AD among *APOE4* carriers.

## 2. METHODS

### 2.1. Study Sample

The Sleep Heart Health Study (SHHS) is a multi-center, community-based cohort study designed to investigate relationships between sleep disorders and health outcomes in U.S. adults. Details of the study design and data collection protocols have been described previously. ^25^ Briefly, between 1995 and 1998, the SHHS enrolled 6,441 adults aged 40 years and older from several established parent cohorts, including the Atherosclerosis Risk in the Community Study (ARIC), ^26^ the Cardiovascular Health Study (CHS), ^27^ the Framingham Heart Study (FHS), ^28^ and a few others. The enrolled participants subsequently completed a single night of unattended, in-home overnight polysomnography. ^25^ Blood samples were collected by the parent cohorts, facilitating *APOE* genotyping.

For the current study, we included participants who completed overnight polysomnography and had available *APOE* genotype data. Because *APOE* genotype was publicly available in only a subset of the parent cohorts (i.e., ARIC, CHS, and FHS), our analytic sample was restricted to participants recruited from these three parent cohorts. The final analytic sample included 3,131 participants between ages 50 and 85 after excluding individuals with missing data. See Supplementary Figure 1 for processes used to derive the analytic sample. All study procedures were approved by the Yale University Institutional Review Board.

### 2.2. Outcomes

Full-montage in-home polysomnography was performed using the Compumedics PS system (Abbotsford, Victoria, Australia) following standardized protocols, including electroencephalography, electrooculography, chin electromyography, thoracic and abdominal plethysmography, airflow measurement, pulse oximetry, electrocardiography, and body position monitoring. ^25^ Sleep stages and arousals were manually scored in 30-second epochs according to American Academy of Sleep Medicine criteria. ^25^ Detailed procedures for the polysomnography, data processing, and sleep scoring have been described previously. ^25^

The outcomes of interest included macro and micro sleep architecture characteristics derived from the overnight in-home polysomnography data. Macro-architecture variables included the proportion of rapid eye movement sleep (%REM) and the proportions of non-REM stages (%N1, %N2, and %N3 or slow-wave sleep [%SWS]), as well as the arousal index (the number of arousals per hour). Percent time spent in each sleep stage was calculated by dividing the total time spent in each stage by total sleep time, providing normalization across individuals with differing sleep durations. Arousal index was computed as the number of arousal events per hour of total sleep time. Microarchitecture variables included N2 spindle density, spindle power, and spindle frequency, along with the average odds ratio product (ORP), a continuous measure of sleep depth, defined as the likelihood of being awake, ranging from 0 (always asleep) to 2.5 (always awake). ^29^ Spindles during N2 were detected using a previously described method based on the Michele Sleep Scoring system. ^30^ Using the detected spindle events, spindle density was defined as the number of valid spindles per minute of N2 sleep. ^31^ Spindle power was defined as the average peak power across all valid spindles, and spindle frequency was defined as the EEG frequency at which peak power occurred, averaged across spindles. ^31^ The ORP was calculated using established methods, ^29,32^ in which the EEG power spectrum in the delta, theta, alpha, and beta bands for each 3 second epoch was used to derive a ratio that reflects the probability of wakefulness based on a large reference dataset. ^29,32^ The resulting ORP values provide a continuous measure of sleep depth, defined as the odds of being awake, which is independent of conventional stage scoring. ^29,32^

### 2.3. Exposure

*APOE* carrier status was categorized into the following four groups based on genotyping performed using DNA extracted from whole blood: *APOE4* heterozygotes (ε2/ε4 or ε3/ε4), *APOE4* homozygotes (ε4/ε4), *APOE2* carriers (ε2/ε2 or ε2/ε3), and *APOE3* homozygotes (ε3/ε3). *APOE4* heterozygotes and homozygotes were analyzed as distinct groups to assess potential dose-dependent effects of *APOE4*. *APOE3* homozygotes served as the reference group as they are the population majority. ^23^ Detailed protocols for blood collection, DNA isolation, and *APOE* genotyping have been described previously. ^33–35^

### 2.4. Statistical Analysis

Differences in sample characteristics by *APOE* genotype were assessed using ANOVA for continuous variables and chi-square for categorical variables. We then assessed the association between *APOE* genotype and each sleep variable using linear regression models, adjusting for age, sex, race, marital status, educational attainment, parent cohort, as well as the frequency of taking sleeping pills (ranging from never to almost always). We did not adjust for medical history, as these factors may mediate or be associated with both *APOE* genotype and sleep, thereby obscuring the association of interest. This approach is appropriate for the proposed analyses, as our goal is to examine genotype-specific sleep characteristics, regardless of whether they are attributable to underlying medical conditions. Analyses were conducted in the full sample and stratified by sex to evaluate potential differences between males and females. Moreover, to test whether age moderated the relationship between *APOE* genotype and sleep architecture, we introduced interactions between *APOE* genotype and age, using both linear and quadratic terms. Age was mean centered using the sample mean to improve applicability of regression estimates to the age range examined in the current study. If an age by APOE genotype interaction was significantly associated with a sleep characteristic, marginal differences between genotypes were calculated across the examined age span to identify the ages at which the given sleep characteristic began to diverge. Statistical significance was defined at the level of *p* < 0.05. All analyses were conducted using Python 3.10 (Python Software Foundation, Wilmington, DE).

Posthoc analyses examined additional sleep characteristics, including wake after sleep onset (WASO; total time spent awake between sleep onset and offset), average wake period per arousal (WASO divided by the total number of arousals), changes in ORP overnight (difference between ORPs of the first and the last hour of sleep), ^36^ and the rate of sleep recovery following arousals (average ORP of 9 seconds following the end of an arousal). ^32,37^

## 3. RESULTS

Table 1 shows the characteristics of the analytic sample. The mean age was 66.7 years (standard deviation=8.8), 52.7% were female, and 94.9% were White. Eighty three percent were married and 35.1% completed ≥16 years of education. Average proportions of REM sleep (%REM) and SWS (%SWS) were 19.9% and 18.6%, respectively. Individuals had a mean of 19.5 arousals per hour. A total of 748 persons (23.9 %) carried one *APOE4* allele and 67 (2.1%) carried two *APOE4* alleles. All sample characteristics were comparable across *APOE* genotype, except for race, where the *APOE2* carriers included lower proportion of individuals who identified as White, and ORP, which was lowest in *APOE4* homozygotes.

**Table 1.**
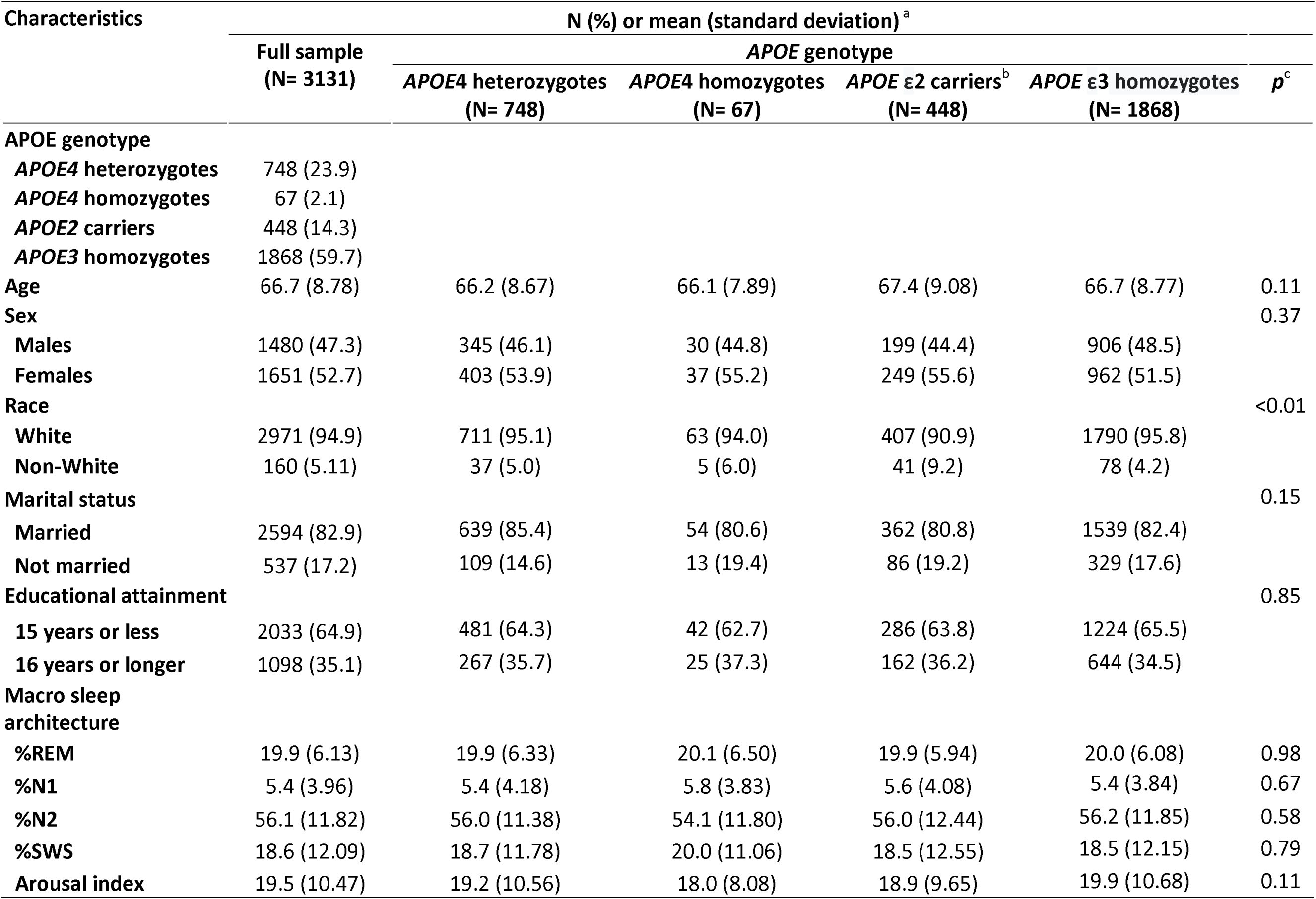
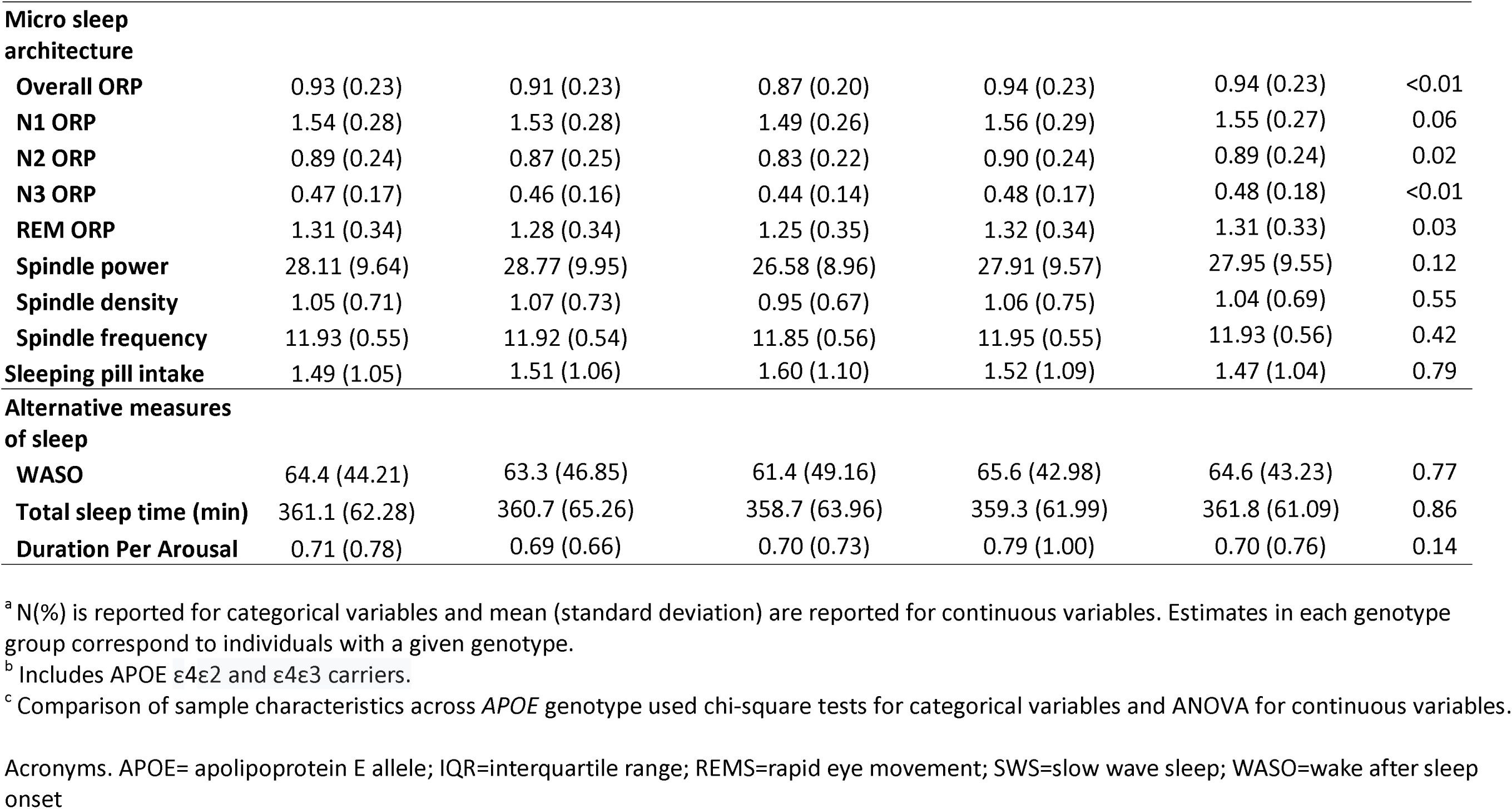
Sample characteristics.

| Characteristics | N (%) or mean (standard deviation) <sup>a</sup> |  |  |  |  | *p* <sup>c</sup> |
| --- | --- | --- | --- | --- | --- | --- |
|  | Full sample<br>(N= 3131) | *APOE* genotype |  |  |  |  |
|  |  | *APOE4* heterozygotes<br>(N= 748) | *APOE4* homozygotes<br>(N= 67) | *APOE* ε2 carriers <sup>b</sup><br>(N= 448) | *APOE* ε3 homozygotes<br>(N= 1868) |  |
| ***APOE* genotype** |  |  |  |  |  |  |
| *APOE4* heterozygotes | 748 (23.9) |  |  |  |  |  |
| *APOE4* homozygotes | 67 (2.1) |  |  |  |  |  |
| *APOE2* carriers | 448 (14.3) |  |  |  |  |  |
| *APOE3* homozygotes | 1868 (59.7) |  |  |  |  |  |
| Age | 66.7 (8.78) | 66.2 (8.67) | 66.1 (7.89) | 67.4 (9.08) | 66.7 (8.77) | 0.11 |
| Sex |  |  |  |  |  | 0.37 |
| Males | 1480 (47.3) | 345 (46.1) | 30 (44.8) | 199 (44.4) | 906 (48.5) |  |
| Females | 1651 (52.7) | 403 (53.9) | 37 (55.2) | 249 (55.6) | 962 (51.5) |  |
| Race |  |  |  |  |  | <0.01 |
| White | 2971 (94.9) | 711 (95.1) | 63 (94.0) | 407 (90.9) | 1790 (95.8) |  |
| Non-White | 160 (5.11) | 37 (5.0) | 5 (6.0) | 41 (9.2) | 78 (4.2) |  |
| Marital status |  |  |  |  |  | 0.15 |
| Married | 2594 (82.9) | 639 (85.4) | 54 (80.6) | 362 (80.8) | 1539 (82.4) |  |
| Not married | 537 (17.2) | 109 (14.6) | 13 (19.4) | 86 (19.2) | 329 (17.6) |  |
| Educational attainment |  |  |  |  |  | 0.85 |
| 15 years or less | 2033 (64.9) | 481 (64.3) | 42 (62.7) | 286 (63.8) | 1224 (65.5) |  |
| 16 years or longer | 1098 (35.1) | 267 (35.7) | 25 (37.3) | 162 (36.2) | 644 (34.5) |  |
| Macro sleep architecture |  |  |  |  |  |  |
| %REM | 19.9 (6.13) | 19.9 (6.33) | 20.1 (6.50) | 19.9 (5.94) | 20.0 (6.08) | 0.98 |
| %N1 | 5.4 (3.96) | 5.4 (4.18) | 5.8 (3.83) | 5.6 (4.08) | 5.4 (3.84) | 0.67 |
| %N2 | 56.1 (11.82) | 56.0 (11.38) | 54.1 (11.80) | 56.0 (12.44) | 56.2 (11.85) | 0.58 |
| %SWS | 18.6 (12.09) | 18.7 (11.78) | 20.0 (11.06) | 18.5 (12.55) | 18.5 (12.15) | 0.79 |
| Arousal index | 19.5 (10.47) | 19.2 (10.56) | 18.0 (8.08) | 18.9 (9.65) | 19.9 (10.68) | 0.11 |
| **Micro sleep architecture** |  |  |  |  |  |  |
| **Overall ORP** | 0.93 (0.23) | 0.91 (0.23) | 0.87 (0.20) | 0.94 (0.23) | 0.94 (0.23) | <0.01 |
| **N1 ORP** | 1.54 (0.28) | 1.53 (0.28) | 1.49 (0.26) | 1.56 (0.29) | 1.55 (0.27) | 0.06 |
| **N2 ORP** | 0.89 (0.24) | 0.87 (0.25) | 0.83 (0.22) | 0.90 (0.24) | 0.89 (0.24) | 0.02 |
| **N3 ORP** | 0.47 (0.17) | 0.46 (0.16) | 0.44 (0.14) | 0.48 (0.17) | 0.48 (0.18) | <0.01 |
| **REM ORP** | 1.31 (0.34) | 1.28 (0.34) | 1.25 (0.35) | 1.32 (0.34) | 1.31 (0.33) | 0.03 |
| **Spindle power** | 28.11 (9.64) | 28.77 (9.95) | 26.58 (8.96) | 27.91 (9.57) | 27.95 (9.55) | 0.12 |
| **Spindle density** | 1.05 (0.71) | 1.07 (0.73) | 0.95 (0.67) | 1.06 (0.75) | 1.04 (0.69) | 0.55 |
| **Spindle frequency** | 11.93 (0.55) | 11.92 (0.54) | 11.85 (0.56) | 11.95 (0.55) | 11.93 (0.56) | 0.42 |
| **Sleeping pill intake** | 1.49 (1.05) | 1.51 (1.06) | 1.60 (1.10) | 1.52 (1.09) | 1.47 (1.04) | 0.79 |
| **Alternative measures of sleep** |  |  |  |  |  |  |
| **WASO** | 64.4 (44.21) | 63.3 (46.85) | 61.4 (49.16) | 65.6 (42.98) | 64.6 (43.23) | 0.77 |
| **Total sleep time (min)** | 361.1 (62.28) | 360.7 (65.26) | 358.7 (63.96) | 359.3 (61.99) | 361.8 (61.09) | 0.86 |
| **Duration Per Arousal** | 0.71 (0.78) | 0.69 (0.66) | 0.70 (0.73) | 0.79 (1.00) | 0.70 (0.76) | 0.14 |

<sup>a</sup> N(%) is reported for categorical variables and mean (standard deviation) are reported for continuous variables. Estimates in each genotype group correspond to individuals with a given genotype.
<sup>b</sup> Includes APOE ε4ε2 and ε4ε3 carriers.
<sup>c</sup> Comparison of sample characteristics across *APOE* genotype used chi-square tests for categorical variables and ANOVA for continuous variables.
Acronyms. APOE= apolipoprotein E allele; IQR=interquartile range; REMS=rapid eye movement; SWS=slow wave sleep; WASO=wake after sleep onset

Table 2 presents associations between macro sleep architecture and *APOE* genotype, estimated using models adjusted for all covariates. In models without interaction terms, %REM, %N1, %N2, %SWS, and the arousal index did not differ significantly across the genotypes. In models that included an interaction between age and *APOE* genotype, the association between genotype and arousal index varied with age, such that *APOE4* homozygotes showed 0.33 fewer arousals per hour per year of age compared to *APOE3* homozygotes (p = 0.04). As a result, *APOE4* homozygotes exhibited significantly fewer arousals than *APOE3* homozygotes beginning at age 70 (Figure 1A). The association of %REM, %N1, %N2, and %SWS with *APOE* genotype did not vary with age. Interaction terms involving quadratic age were not significant in any model, and therefore were excluded.

**Figure 1.**
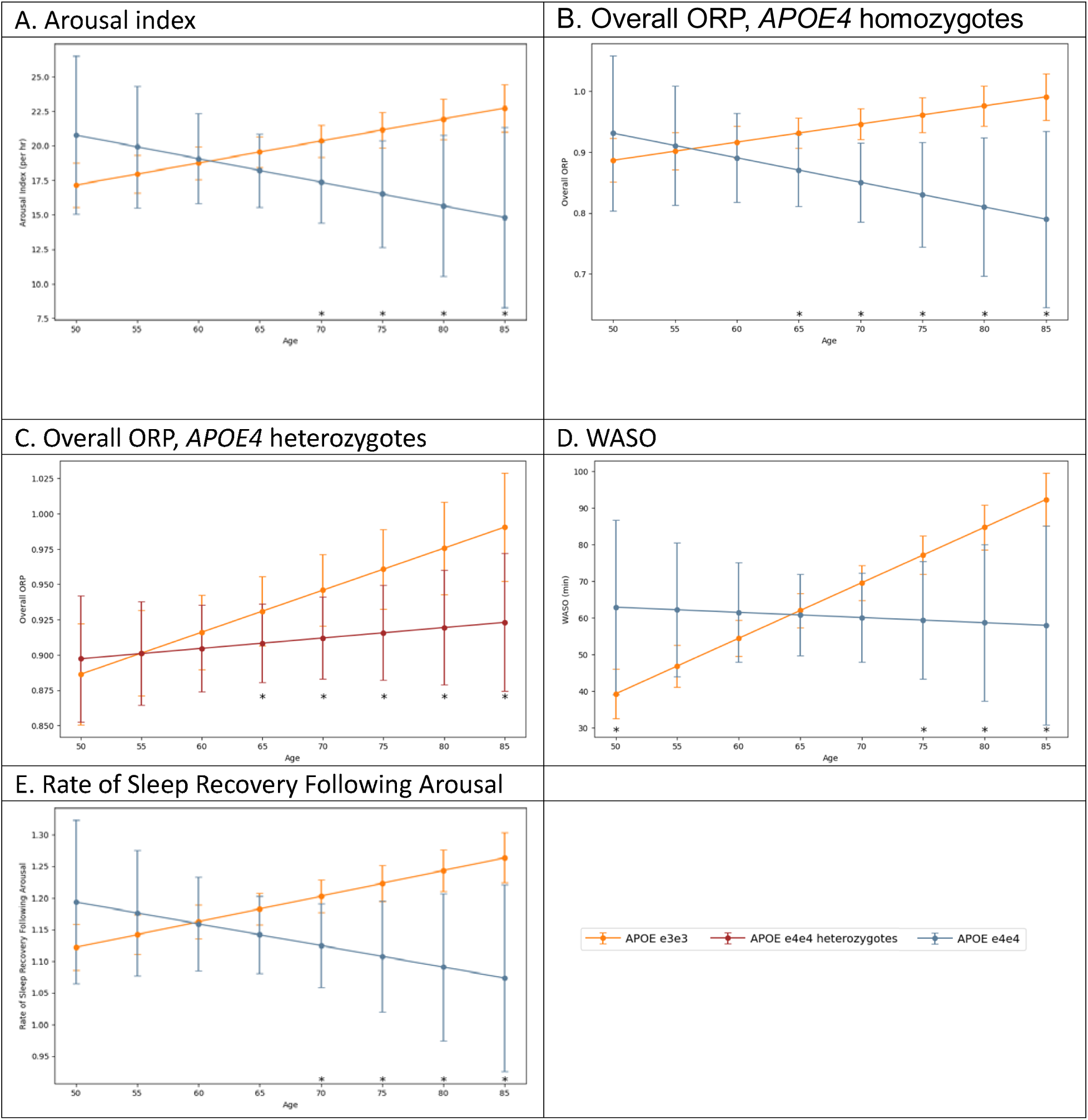
Marginal differences in arousal characteristics by APOE genotype and age. Acronyms. ORP=odds ratio product; WASO=wake after sleep onset

**Table 2.**
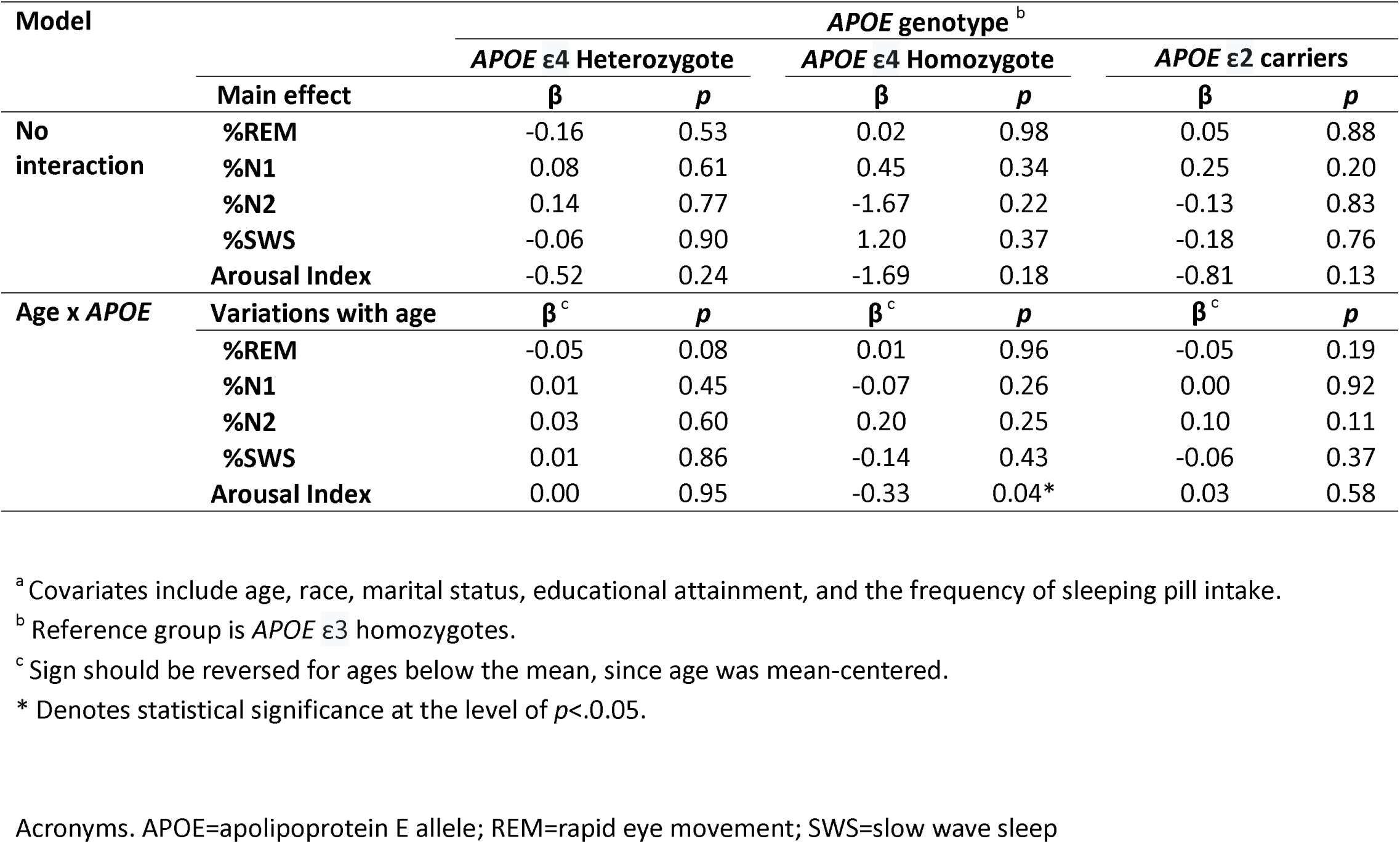
Association between APOE genotype and sleep architecture.

Table 3 presents associations between sleep microarchitecture and *APOE* genotype, estimated using models adjusted for all covariates. In models without interaction terms, differences in spindle characteristics did not reach statistical significance, although *APOE4* homozygotes exhibited a trend towards lower spindle power (p=0.059) and density (p=0.069) compared to *APOE3* homozygotes. Moreover, ORP decreased in a dose-response pattern with each additional *APOE4* allele, with both *APOE4* homozygotes and heterozygotes showing significantly lower ORP than *APOE3* homozygotes during overall, N2, and N3 sleep (β for overall sleep, APOE4 homozygote = −0.07, p = 0.01). The lowest mean levels of ORP were consistently found in *APOE4* homozygotes across all sleep stages. In addition, *APOE4* heterozygotes showed significantly lower ORP than *APOE3* homozygotes across all sleep stages (ps < 0.05). In models that additionally included an interaction between age and *APOE* genotype, the difference in ORP between *APOE4* carriers and *APOE3* homozygotes widened with advancing age for overall sleep and multiple stages including REM. Figures 1B and 1C graphically demonstrate that *APOE4* homozygotes and heterozygotes exhibit a steeper downward trend in ORP than *APOE3* homozygotes between ages 50 and 85 for overall sleep. *APOE2* carriers did not differ from *APOE3* homozygotes in ORP across the age span examined in this study.

**Table 3.** Association between APOE genotype and micro sleep architecture (post-hoc analyses)

| Model |  | *APOE4* genotype<sup>b</sup> |  |  |  |  |  |
| --- | --- | --- | --- | --- | --- | --- | --- |
|  |  | *APOE* ε4 Heterozygote |  | *APOE* ε4 Homozygote |  | *APOE* 2 carriers |  |
|  |  | β | *p* | β | *p* | β | *p* |
| **No interaction** | **Spindle power** | 0.436 | 0.244 | -2.028 | 0.059 | -0.025 | 0.957 |
|  | **Spindle density** | -0.005 | 0.860 | -0.144 | 0.069 | 0.031 | 0.355 |
|  | **Spindle frequency** | -0.037 | 0.099 | -0.107 | 0.093 | 0.030 | 0.271 |
|  | **Overall ORP** | -0.026 | 0.009* | -0.069 | 0.014* | -0.001 | 0.906 |
|  | **N1 ORP** | -0.024 | 0.042* | -0.059 | 0.084 | 0.000 | 0.991 |
|  | **N2 ORP** | -0.025 | 0.018* | -0.065 | 0.030* | -0.005 | 0.687 |
|  | **N3 ORP** | -0.021 | 0.005* | -0.047 | 0.032* | -0.005 | 0.612 |
|  | **REM ORP** | -0.033 | 0.020* | -0.073 | 0.080 | 0.000 | 0.991 |
| **Age x *APOE*** | **Variations with age** | β<sup>c</sup> | *p* | β<sup>c</sup> | *p* | β<sup>c</sup> | *P* |
|  | **Spindle power** | 0.018 | 0.675 | -0.046 | 0.739 | -0.014 | 0.788 |
|  | **Spindle density** | -0.003 | 0.316 | -0.004 | 0.688 | 0.001 | 0.831 |
|  | **Spindle frequency** | 0.001 | 0.754 | -0.011 | 0.182 | 0.001 | 0.808 |
|  | **Overall ORP** | -0.002 | 0.047* | -0.007 | 0.050 | 0.000 | 0.833 |
|  | **N1 ORP** | -0.003 | 0.040* | -0.005 | 0.241 | 0.002 | 0.210 |
|  | **N2 ORP** | -0.002 | 0.139 | -0.008 | 0.026* | 0.000 | 0.852 |
|  | **N3 ORP** | 0.000 | 0.624 | -0.003 | 0.302 | -0.001 | 0.366 |
|  | **REM ORP** | -0.004 | 0.025* | -0.009 | 0.080 | 0.000 | 0.901 |

<sup>a</sup> Covariates include age, race, marital status, educational attainment, and the frequency of sleeping pill intake.<sup>b</sup> Reference group is *APOE* ε3 homozygotes.<sup>c</sup> Sign should be reversed for ages below the mean, since age was mean-centered.

Table 4 presents findings from post hoc analyses that examined additional arousal-related characteristics - including WASO, average arousal duration, overnight change in ORP, and the rate of sleep recovery - to further characterize the observed decrease in arousal index and ORP among *APOE4* carriers. Consistent with the main analyses, additional arousal-related characteristics did not differ significantly across *APOE* genotypes in models without an interaction term. However, *APOE4* homozygotes experienced 1.7 fewer minutes of WASO per year of age relative to *APOE3* homozygotes (p = 0.01), leading to WASO remaining relatively stable across the examined age range in *APOE4* homozygotes, whereas it increased monotonically in *APOE3* homozygotes (please see Figure 1D). Thus, both groups exhibited approximately 60 minutes of WASO at age 65, but by age 75, *APOE4* homozygotes showed substantially shorter WASO. The rate of sleep recovery following arousals mirrors the findings on WASO (Figure 1E). No genotype-related differences were observed in average wake period per arousal and overnight change in ORP.

**Table 4.** Association between APOE genotype and alternative measures of sleep (post-hoc analyses)

| Model |  | *APOE4* genotype<sup>b</sup> |  |  |  |  |  |
| --- | --- | --- | --- | --- | --- | --- | --- |
|  |  | *APOE* ε4 Heterozygote |  | *APOE* ε4 Homozygote |  | *APOE* 2 carriers |  |
|  |  | β | p | β | p | β | p |
| **No interaction** | **WASO** | -0.691 | 0.707 | -2.983 | 0.572 | 0.077 | 0.972 |
|  | **Average wake period per arousals** | -0.006 | 0.869 | -0.007 | 0.943 | 0.065 | 0.105 |
|  | **Change in ORP over night** | -0.007 | 0.517 | 0.009 | 0.765 | 0.015 | 0.256 |
|  | **Rate of sleep recovery following arousals** | -0.010 | 0.314 | -0.049 | 0.089 | 0.000 | 0.978 |
| **Age x *APOE*** | **Variations with age** | β<sup>c</sup> | p | β<sup>c</sup> | p | β<sup>c</sup> | p |
|  | **WASO** | 0.117 | 0.579 | -1.656 | 0.013* | -0.064 | 0.795 |
|  | **Average wake period per arousals** | 0.001 | 0.711 | -0.013 | 0.279 | -0.004 | 0.412 |
|  | **Change in ORP over night** | 0.000 | 0.895 | -0.005 | 0.200 | 0.002 | 0.316 |
|  | **Rate of sleep recovery following arousals** | -0.001 | 0.216 | -0.008 | 0.040* | -0.002 | 0.074 |

Acronyms. REMS=rapid eye movement; SWS=slow wave sleep; WASO=wake after sleep onset
<sup>a</sup> Covariates include age, race, marital status, educational attainment, and the frequency of sleeping pill intake.
<sup>b</sup> Reference group is *APOE* ε3 homozygotes.
<sup>c</sup> Sign should be reversed for ages below the mean, since age was mean-centered.
\* Denotes statistical significance at the level of $p < 0.05$ .

Supplementary Tables 1 and 2 present estimates from sex-stratified models adjusted for all covariates. Overall, the findings in men and women were consistent with those observed in the full sample. Significance levels were slightly attenuated, particularly among women, due to smaller sample sizes and reduced representation of *APOE4* homozygotes due to stratification.

## 4. DISCUSSION

The primary motivation of the current study was to investigate whether sleep architecture may serve as an underlying pathway linking *APOE4* alleles to elevated AD risk, through an examination of both macro and microarchitecture of sleep. We had hypothesized that *APOE4* carriers would show alterations in sleep architecture that are robustly associated with neurodegeneration, including greater sleep fragmentation, less SWS, and fewer spindles. ^13^ Contrary to this hypothesis, we found consistent indications that *APOE4* carriers, particularly *APOE4* homozygotes, are less likely to experience sleep fragmentation. The analyses of sleep microarchitecture showed overall ORP - a measure of sleep depth where lower values represent deeper sleep - to decrease in a dose response manner with each *APOE4* allele. *APOE4* homozygotes exhibited the lowest ORP across genotypes, indicating having the deepest sleep, defined as the lowest likelihood of being awake given an EEG profile. ^29,32,36–38^ The gap in ORP between *APOE4* carriers and *APOE3* homozygotes widened with age. Notably, *APOE4* homozygotes showed an age-related increase in sleep depth, whereas *APOE3* homozygotes showed a reduction in sleep depth with age. These findings were further supported by the analysis of arousal index, WASO, sleep recovery rate following an arousal - defined as average ORP during 9 seconds following an arousal where lower values indicate faster recovery. All three characteristics exhibited age-related declines in *APOE4* homozygotes consistent with those observed for ORP. Taken together, *APOE4* carriers, especially *APOE4* homozygotes, may have an elevated thresholds for arousals and awakenings. As a result, they show decreases in sleep fragmentation with age, an opposite trend observed among *APOE3* homozygotes, whose levels of sleep fragmentation increases with age, a pattern that has been documented in normal aging.

At first glance, these findings seem to suggest that *APOE4* homozygotes experience less sleep fragmentation, which is often equated with “healthy sleep”. However, interpreting elevated arousal thresholds in *APOE4* carriers requires caution, when considering that *APOE4* homozygotes were found to also exhibit less spindle activity compared to *APOE3* homozygotes. In particular*, APOE4* homozygotes exhibited trends towards reduced spindle density and power, patterns that have been robustly associated with AD and cognitive aging. ^10–12,39,40^ While the exact function of sleep spindles still remains unclear, evidence thus far has shown it to play key roles in hippocampal-cortical cross-talk necessary for memory consolidation. ^41^ Furthermore, a recent study in neurofluids showed that sleep spindles are temporally linked to brain waste clearance, directly preceding microarousals that facilitate cerebrospinal movement that presumably facilitates solute transport and removal from the brain. ^9^ Our findings contribute to this line of literature on sleep spindles and brain health by showing that *APOE4* homozygotes, who have the highest genetic risk of AD, may experience alterations in spindle activity during middle and late adulthood, which may contribute to AD progression by precluding its propitious effects. To the best of our knowledge, the current study is the first to examine differences in spindle activity by *APOE* genotype.

Furthermore, it is important to highlight that lower levels of sleep fragmentation found in *APOE4* carriers were not accompanied by increases in REM sleep or SWS. Thus, these findings support the view that the observed elevation in arousal threshold among *APOE4* carriers may not indicate more restorative sleep, ^2,3,13,42,43^ but may instead be associated with neurodegeneration in regions that regulate sleep and wakefulness, such as the locus coeruleus and basal forebrain. Such neurodegenerative changes may blunt cortical activation required for arousals and awakenings. ^5–8,44^ While arousals are often regarded as detrimental to health, emerging evidence suggests that sleep micro-fragmentation or microarousals – defined as very short high frequency shift in EEG that do not lead to an awakening - may benefit brain health. ^45^ For example, microarousals have been shown to enhance brain waste clearance by promoting large-amplitude cerebral vasomotion and cerebrospinal fluid flow, facilitating solute transport and removal from the brain. ^9^ In addition, more frequent micro fragmentation of sleep has been associated with lower Aβ burden. ^46^ Thus, it is possible that *APOE4* homozygotes experience fewer of these potentially beneficial microarousals due to elevated arousal thresholds, although we did not find between-genotype differences in average wake period per arousal. Additional empirical research is needed to characterize arousal features in *APOE4* carriers, including distinctions across arousal subtypes described in prior research, ^45^ and to clarify how these features may mediate the risk of AD.

To date, two studies have examined the association between *APOE* genotype and sleep macro architecture, reporting heterogeneous findings after examining different samples. Tranah et al. examined community-dwelling older men (mean age=77 years; N=2,302, including 515 *APOE4* heterozygotes and 40 homozygotes) and found that *APOE4* homozygotes had longer SWS and shorter WASO compared to non-carriers. ^16^ Our findings are partially consistent with those of Tranah et al. ^16^ as we found *APOE4* homozygotes to have elevated arousal and awakening thresholds, as shown by multiple metrics including WASO. However, we did not find *APOE4* carriers to have longer SWS. Reasons behind this discrepancy in findings is unclear, but may be potentially due to our operational definition of SWS as its proportion out of total sleep time, which can be different from its absolute duration used by Tranah et al. In comparison, Andre et al. examined a smaller, mixed-sex sample (mean age=69 years; N = 198, including 41 heterozygotes and 5 homozygotes) and reported that *APOE4* carriers (without distinguishing between heterozygotes and homozygotes) had shorter REM sleep, both in absolute duration and as a proportion of total sleep time. ^17^ They did not find differences in SWS and WASO reported by Tranah et al. ^17^ Our findings also largely diverge from those of Andre et al., as we did not find differences in REM sleep by *APOE* genotype. These differences may reflect the larger number of *APOE4* heterozygotes and homozygotes included in our study, which allowed us to distinguish between the two genotypes. Collectively, the majority of studies to date – which include the current study and Tranah et al. - suggests that *APOE4* homozygotes may experience less sleep fragmentation in late adulthood. Further validation studies are required to clarify the impact of *APOE* genotype on REM and SWS, in a sample with sufficient representation of *APOE4* homozygotes.

The strengths of the current study include the following. First, this study is the first to examine sleep microarchitecture, including ORP and sleep spindles, allowing for the detection of micro-time-scale differences in sleep quality. Second, this study used gold-standard methods to comprehensively assess the features of sleep macro- and micro-architecture that have been previously associated with Alzheimer’s disease risk. ^14,47^ Third, this study included the largest number of *APOE4* heterozygotes and homozygotes among all existing studies on this topic, while focusing on community-dwelling adults, enhancing the applicability and generalizability of the findings. Fourth, the inclusion of middle-aged adults allowed for the examination of age-related variations in the association between *APOE* genotype and sleep. Age related differences in arousal and awakening thresholds reported in this study may be important for clinical practice in understanding how sleep vulnerability evolve across the lifespan and how these patterns may inform early identification of individuals at elevated risk for AD.

Nonetheless, this study is not without limitations. First, polysomnography does not capture all relevant features of sleep health. For instance, daytime napping behavior, which is a sign of non-restorative sleep and a risk factor of AD, ^48^ may be relevant. Second, we did not examine microstructural features of sleep architecture other than ORP or sleep spindles, such delta wave density. Third, sleep quality was assessed based on a single night of polysomnography, which may be influenced by the “first-night effect”, a well-documented phenomenon wherein sleep is disturbed during the initial night of monitoring. ^49^ Thus, our estimates of sleep architecture may not reflect individuals’ habitual sleep patterns. Fourth, over 90% of the analytic sample identified as White, decreasing generalizability of our results to other racial-ethnic groups. To address these limitations, future studies should examine the association between e *APOE* genotype and sleep using more comprehensive assessments of sleep, including daytime napping, applied to more diverse populations.

## Conclusions

This study investigated the association of *APOE* genotype with the macro and microarchitecture of sleep in a large population based sample of middle aged and older adults. Contrary to expectations, robust evidence was found suggesting that *APOE4* carriers, particularly *APOE4* homozygotes, experience less sleep fragmentation than *APOE3* homozygotes, the population majority with lower AD risk. Notably, *APOE4* heterozygotes and homozygotes had deeper sleep on average, and the carriers’ sleep depth increased further with advancing age. In addition, *APOE4* homozygotes exhibited fewer arousals, less WASO, and faster sleep recovery after arousals with advancing age. Nonetheless, lower levels of sleep fragmentation observed in *APOE4* carriers were not accompanied by other indicators of healthy sleep. In particular, *APOE4* homozygotes showed a trend towards reduced spindle activity, including spindle density and power. Also, both *APOE4* homozygotes and heterozygotes exhibited comparable sleep macro architecture as *APOE3* homozygotes. These findings collectively suggest that elevated thresholds for arousals and awakenings observed in *APOE4* carriers may reflect blunted cortical activation due to neurodegeneration in regions that regulate sleep wake transitions rather than healthier sleep. Future research is needed to better characterize sleep fragmentation among *APOE4* carriers, with a focus on the prevalence of different arousal subtypes and how each type is associated with AD risk.

## ACKNOWLEDGEMENTS

This Manuscript was prepared using SHHS Research Materials obtained from the NHLBI Biologic Specimen and Data Repository Information Coordinating Center and the National Sleep Research Resource, does not necessarily reflect the opinions or views of the SHHS, NSRR, or the NHLBI. The Sleep Heart Health Study (SHHS) was supported by National Heart, Lung, and Blood Institute cooperative agreements U01HL53916 (University of California, Davis), U01HL53931 (New York University), U01HL53934 (University of Minnesota), U01HL53937 and U01HL64360 (Johns Hopkins University), U01HL53938 (University of Arizona), U01HL53940 (University of Washington), U01HL53941 (Boston University), and U01HL63463 (Case Western Reserve University). The National Sleep Research Resource was supported by the National Heart, Lung, and Blood Institute (R24 HL114473, 75N92019R002).

## FUNDING

This work was supported by the National Institute of Health (P30AG021342) and the Alzheimer’s Association (AARFD-24-1306796).

This work was supported by the National Institutes of Health under award number [P30AG021342]. This manuscript is the result of funding in whole or in part by the National Institutes of Health (NIH). It is subject to the NIH Public Access Policy. Through acceptance of this federal funding, NIH has been given a right to make this manuscript publicly available in PubMed Central upon the Official Date of Publication, as defined by NIH.

## DISCOLSURES

Outside of the current work, Orfeu M. Buxton discloses that he received two subcontract grants to Penn State from Mobile Sleep Technologies (NSF/STTR #1622766, NIH/NIA SBIR R43-AG056250, R44-AG056250), received honoraria/travel support for lectures from New York University, Tufts School of Dental Medicine, University of Utah, University of Arizona, and University of Miami; consulting fees from Georgia State University and Harvard Chan School of Public Health; and receives an honorarium for his role as the Editor in Chief of Sleep Health sleephealthjournal.org.

Gawon Cho, Anne Chen, Eunyoung Choi, Daniel Kay, and Brienne Miner do not have financial support to declare and report absence of conflict of interest.

The initial draft has been posted as a preprint on Biorxiv: 10.1101/2025.05.30.656615

## Notes

https://biolincc.nhlbi.nih.gov/studies/shhs/

https://biolincc.nhlbi.nih.gov/studies/aric/

https://biolincc.nhlbi.nih.gov/studies/chs/

https://www.ncbi.nlm.nih.gov/projects/gap/cgi-bin/study.cgi?study_id=phs000007.v33.p14

### Summary of Updates

Analysis was broadened to the investigation of micro sleep architecture.

