## Supplementary Table for "Comprehensive investigation of sleep architecture by *APOE* genotype: *APOE4* homozygotes have higher arousal thresholds"

**Supplementary Figure 1.** Identification of analytic sample

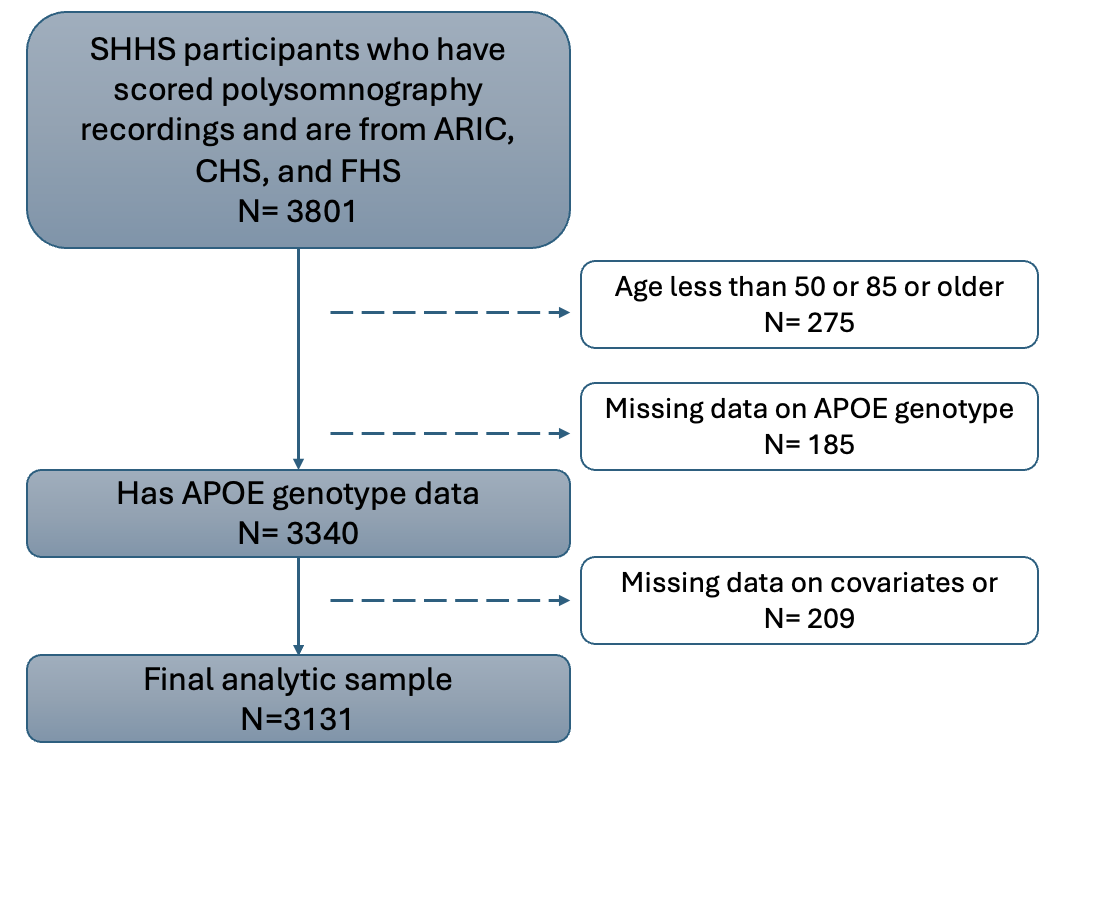

**Supplementary Table 1.** Association between *APOE* genotype and sleep architecture stratified by sex ^a^

| **Model** | | ***APOE* genotype** ^b^ | | | | | | | |
| --- | --- | --- | --- | --- | --- | --- | --- | --- | --- |
|  |  | ***APOE*4 Heterozygote** | |  | ***APOE*4 Homozygote** | |  | ***APOE*2 carriers** | |
|  | **Sleep architecture** | **β** | ***p*** |  | **β** | ***p*** |  | **β** | ***p*** |
| **Men**  **n= 1480** | **%REMS** | -0.23 | 0.55 |  | 1.75 | 0.12 |  | 0.17 | 0.72 |
|  | **%N1** | 0.21 | 0.45 |  | 0.34 | 0.68 |  | 0.39 | 0.26 |
|  | **%N2** | -0.81 | 0.21 |  | -3.95 | 0.04* |  | -1.24 | 0.12 |
|  | **%SWS** | 0.83 | 0.20 |  | 1.86 | 0.33 |  | 0.68 | 0.39 |
|  | **Arousal index** | -0.02 | 0.98 |  | -1.82 | 0.39 |  | -1.83 | 0.04* |
|  | **Spindle power** | 0.44 | 0.37 |  | -3.19 | 0.03* |  | -0.24 | 0.70 |
|  | **Spindle density** | -0.04 | 0.27 |  | -0.20 | 0.06 |  | 0.01 | 0.90 |
|  | **Spindle frequency** | -0.05 | 0.13 |  | -0.10 | 0.30 |  | 0.02 | 0.66 |
|  | **Overall ORP** | -0.02 | 0.11 |  | -0.10 | 0.02* |  | -0.01 | 0.51 |
|  | **N1 ORP** | -0.02 | 0.20 |  | -0.09 | 0.06 |  | -0.01 | 0.60 |
|  | **N2 ORP** | -0.02 | 0.22 |  | -0.10 | 0.02* |  | -0.01 | 0.55 |
|  | **N3 ORP** | -0.02 | 0.15 |  | -0.05 | 0.14 |  | -0.01 | 0.71 |
|  | **REM ORP** | -0.02 | 0.22 |  | -0.11 | 0.06 |  | -0.02 | 0.41 |
| **Women**  **n= 1651** | **Sleep architecture** | **β** | ***p*** |  | **β** | ***p*** |  | **β** | ***p*** |
|  | **%REMS** | -0.13 | 0.71 |  | -1.40 | 0.16 |  | -0.11 | 0.80 |
|  | **%N1** | -0.08 | 0.64 |  | 0.43 | 0.39 |  | 0.14 | 0.51 |
|  | **%N2** | 0.80 | 0.25 |  | 0.23 | 0.91 |  | 0.88 | 0.29 |
|  | **%SWS** | -0.58 | 0.39 |  | 0.74 | 0.70 |  | -0.91 | 0.26 |
|  | **Arousal index** | -0.94 | 0.08 |  | -1.47 | 0.33 |  | -0.03 | 0.96 |
|  | **Spindle power** | 0.47 | 0.40 |  | -1.07 | 0.49 |  | 0.21 | 0.75 |
|  | **Spindle density** | 0.03 | 0.52 |  | -0.10 | 0.40 |  | 0.06 | 0.25 |
|  | **Spindle frequency** | -0.02 | 0.49 |  | -0.11 | 0.20 |  | 0.04 | 0.26 |
|  | **Overall ORP** | -0.03 | 0.04* |  | -0.05 | 0.20 |  | 0.01 | 0.61 |
|  | **N1 ORP** | -0.03 | 0.12 |  | -0.03 | 0.50 |  | 0.01 | 0.61 |
|  | **N2 ORP** | -0.03 | 0.06 |  | -0.04 | 0.38 |  | 0.00 | 0.97 |
|  | **N3 ORP** | -0.02 | 0.02* |  | -0.05 | 0.10 |  | 0.00 | 0.80 |
|  | **REM ORP** | -0.04 | 0.06 |  | -0.05 | 0.42 |  | 0.02 | 0.40 |

^a^ Covariates include age, race, marital status, educational attainment, and the frequency of sleeping pill intake.

^b^ Reference group is *APOE* ε3 homozygotes.

**Supplementary Table 2.** Association between *APOE* genotype and alternative measures of sleep stratified by sex^a^

| **Model** | | ***APOE4* genotype** ^b^ | | | | | | | |
| --- | --- | --- | --- | --- | --- | --- | --- | --- | --- |
|  | **Other sleep characteristics** | ***APOE4* Heterozygote** | |  | ***APOE4* Homozygote** | |  | ***APOE2* carriers** | |
|  |  | **β** | ***p*** |  | **β** | ***p*** |  | **β** | ***p*** |
| **Men**  **n= 1480** |  |  |  |  |  |  |  |  |  |
|  | **WASO** | -1.26 | 0.66 |  | 2.55 | 0.76 |  | 0.08 | 0.98 |
|  | **Duration Per Arousal** | -0.01 | 0.90 |  | 0.03 | 0.85 |  | 0.13 | 0.05 |
|  | **Change in ORP over night** | -0.01 | 0.50 |  | -0.07 | 0.14 |  | 0.03 | 0.16 |
|  | **Rate of sleep recovery following arousals** | 0.00 | 0.94 |  | -0.09 | 0.03* |  | 0.01 | 0.67 |
| **Women**  **n= 1651** |  | **β** | ***P*** |  | **β** | ***P*** |  | **β** | ***P*** |
|  | **WASO** | -0.73 | 0.76 |  | -7.89 | 0.24 |  | 0.36 | 0.90 |
|  | **Duration Per Arousal** | -0.01 | 0.79 |  | -0.06 | 0.60 |  | 0.02 | 0.63 |
|  | **Change in ORP over night** | 0.00 | 0.76 |  | 0.07 | 0.09 |  | 0.01 | 0.77 |
|  | **Rate of sleep recovery following arousals** | -0.02 | 0.19 |  | -0.02 | 0.65 |  | -0.01 | 0.77 |

^a^ Covariates include age, race, marital status, educational attainment, and the frequency of sleeping pill intake.

^b^ Reference group is *APOE* ε3 homozygotes.
